# Optimizing femtosecond laser fabricating strategy of Nitinol hypotube for vascular interventional surgery

**DOI:** 10.1101/2024.12.01.626210

**Authors:** Yunfeng Song, Zhiwei Yang, Tiangang Xu, Hao Xie, Depeng Wang, Lan Chen, Xusan Yang

**Author notes:** Those authors contribute equally (Yunfeng Song, Zhiwei Yang).

## Abstract

Guidewires and catheters are essential in minimally invasive procedures, particularly in accessing targets within the vascular system for neuron surgery. Femtosecond lasers can precisely cut nickel-titanium micro tubes, known as Nitinol hypotubes, serving as the heads of micro guidewires while protecting their core tips. The quality and geometry of these hypotubes depend on the focus of the laser cutting system. We evaluated the cutting efficiency and quality of Nitinol hypotubes using objective lenses with three numerical apertures (NA). Quantitative analysis indicated that higher NA objectives yield better surface quality, resulting in lower debris. Additionally, cutting with high NA objectives produced significantly lower arithmetic mean surface roughness and enhanced mechanical properties. Using a two-temperature model, we simulated the interaction between femtosecond lasers and Nitinol, showing that higher NA objectives increase electron temperatures, thus improving material removal efficiency. This study also tested the optimized Nitinol hypotube prformance in simulations of invasive neuron surgery.

## Introduction

The nickel-titanium alloy (Nitinol), often employed in the creation of medical equipment, is valued for its inherent characteristics like shape memory effect, superelasticity, compatibility with biological tissues, and resistance to corrosion^1^. Guidewires and catheters are extensively utilized in endovascular minimally invasive procedures, navigating through the circulatory system to reach the intended site, particularly in cardiac and neurosurgical operations^2,3^. Nitinol hypotubes acts as the head of the micro-guidewire and can offer preventive measures against the core tip of the micro-guidewire^4^. With the advancements in laser, computer, and mechanical engineering technologies, the 3D laser cutting process has emerged as the preferred method for the production of Nitinol hypotubes, attributed to its non-contact nature, precision, and focused energy application^5^. Depending on the pulse duration, lasers can be divided into two groups: wide-pulse lasers (with pulse durations >10 ps) and narrow-pulse lasers (with pulse durations <10 ps). Nd: YAG lasers and fiber lasers fall into the wide-pulse category, whereas picosecond and femtosecond lasers are considered ultrafast. Nd: YAG lasers have been the mainstay in fabrication, utilizing a predominantly thermal process for material removal, which is centered on the ejection of melted material^6,7^.Consequently, The advantages of femtosecond lasers include the exceptionally high-quality cut kerf, the narrow kerf width, and the reduction or absence of thermal effects on the material being processed. The downsides are the high cost and the low ablation rate. Therefore, from a financial standpoint, the use of these laser systems is cost-effective only for components that have stringent requirements for cut quality and thermal stress or for extremely thin materials (a few hundred micrometers thick Nitinol micro-tubes), attributed to the brief duration of the laser’s interaction with the material^8^. Nevertheless, a primary challenge in implementing this technology lies in the formation of debris and recast layers, as the surface of the material that has been ablated is often encircled by redeposited vaporized material, which subsequently requires removal through alternative approaches^9^. While these obstacles may result in material properties that are not optimal for medical-grade parts, by carefully choosing the best laser-cutting parameters, the adverse effects of the laser-cutting process can be substantially lessened or alleviated.

In recent years, there have been numerous efforts to optimize the performance of Nitinol materials, particularly in the field of femtosecond laser processing. Research has primarily focused on varying key femtosecond laser parameters, including repetition rate, pulse energy, and laser movement paths, to improve the material’s machining precision, surface quality, and structural integrity during fabrication^10,11^. Studies have demonstrated that adjusting the repetition rate and pulse energy can effectively control the heat-affected zone (HAZ) and material removal rate during laser processing, which is crucial for minimizing thermal damage and enhancing the final product’s performance^12,13^. Recent studies highlight the importance of optimizing laser parameters for achieving high-quality laser processing of Nitinol but fail to delve deeply into the effect of focal size^14^. However, while these studies have provided valuable insights into how these laser parameters influence the material behavior and processing outcomes, one critical parameter that remains underexplored is the focal size of the laser beam. The focal size, which is determined by the numerical aperture (NA) of the focusing optics, plays a crucial role in defining the laser spot size and energy density at the focal point, directly influencing the quality of laser material interaction^12^. For high-precision applications, such as the fabrication of Nitinol hypotubes fabrication, optimizing the NA is essential for achieving clean, precise cuts without causing undesirable thermal effects like cracking or discoloration^15^. Despite its importance, there is limited research specifically addressing the impact of NA on high-resolution Nitinol laser processing, which may limit advancements in micromachining and medical device manufacturing. Further investigations into this aspect could provide insights into improving processing strategies for high-precision Nitinol hypotubes fabrication, ensuring better product reliability and performance in medical applications. This paper presents the influence of NA in Nitinol hypotubes later machining measurements. Here, we first demonstrate experimentally that the differences in focal size caused by NA enable varying degrees of roughness on the lateral and axial surface of the Nitinol hypotubes, Then, the storage modulus of Nitinol hypotubes with the same structure was compared. It was found that hypotubes with high NA exhibited the best overall mechanical performance. Furthermore, we employ finite element simulation to explore the in-depth reasons for these differences, and finally, in vitro experiments were conducted to validate the application effectiveness of Nitinol hypotubes in neuro-interventional guidewires.

## Results

### Debris analysis with different NA

When all other variables are kept constant, the effect of altering the NA from 0.030 to 0.015 is illustrated in Fig.1a and Fig.1d. Employing higher NA, the laser fabrication process resulted in less debris in lateral kerfs. With NA values of 0.019 and 0.015, obvious debris was present on the laser fabrication kerfs, with a debris accumulation width of 7.48 μm and 12.28 μm. Nitinol hypotubes machined with a high NA of 0.030 exhibited the shortest debris accumulation on the surface, with a debris accumulation width of 4.35 μm. Furthermore, Fig.1b and Fig.1e demonstrates that the unobstructed area on the ablation kerf surface of Nitinol hypotube was 8.5% of the total image area for NA=0.030. In the same imaging field, SEM images reveal that the unobstructed area proportions were 8% for NA=0.019 and only about 5.2% for NA=0.015. Moreover, following acid treatment, Fig.1c shows that the kerf edges exhibited an increased presence of jagged microstructures and microcracks with rising NA. In general, it was noted that focusing lenses with a higher NA are more efficient at expelling the debris from the laser fabricating kerf, which results in reduced debris sticking to the surface of the hypotubes.

**Figure 1.**
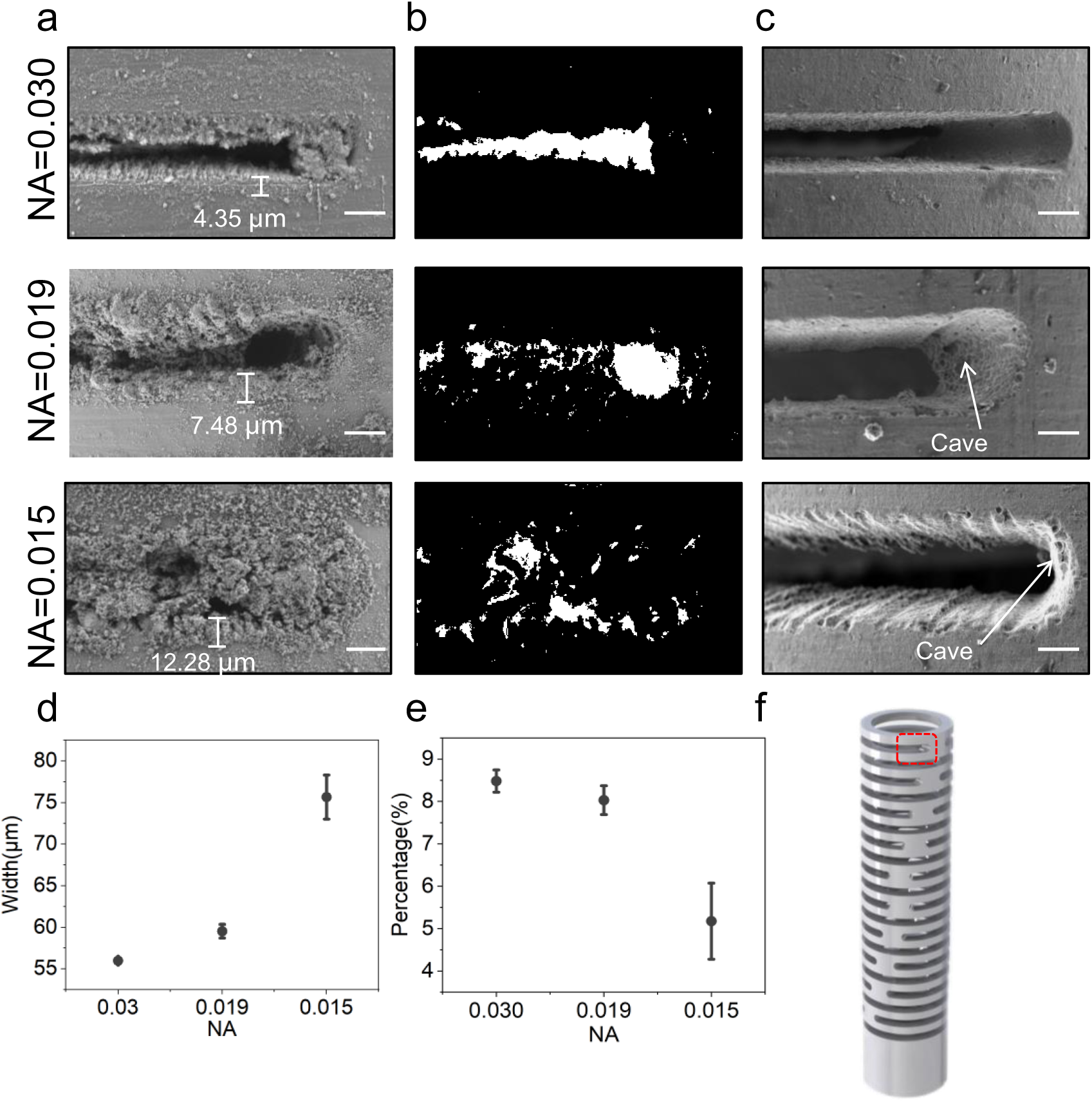
Lateral SEM images and quantitative analysis of the kerfs in Nitinol hypotubes fabricated with femtosecond laser. **a**, Lateral SEM morphology of the fabrication hypotubes before acid dipping, NA=0.030 (top), NA=0.019 (middle), NA=0.015 (bottom). **b**, Binary images of the kerfs in Nitinol hypotubes fabricated with femtosecond laser. **c**, Lateral SEM morphology of the fabrication hypotubes after acid dipping, NA=0.030 (top), NA=0.019 (middle), NA=0.015 (bottom). **d**, Comparative study of experimental methods to assess the width of debris at varying NA, each group of NA is replicated three samples to minimize experimental variability. **e**, Femtosecond laser fabricating debris blockage percentage comparison at various NA, eeach group of NA is replicated three samples to minimize experimental variability. **f**, Schematic diagram of measuring the lateral position. Scale bar, 10 μm.

Additionally, analyzing the axial view of the laser-cut samples can offer further evidence regarding the quantity of debris present on the samples post-fabrication. According to Fig.2a, the maximum accumulated width of the axial debris in the Nitinol hypotube is minimized at NA of 0.030, measuring only 55.94 μm. In comparison, the debris accumulation width at an NA of 0.019 is similar to that at 0.030 NA, measuring 59.91 μm, but the debris accumulation width at an NA of 0.015 is 77.71 μm.

**Figure 2.**
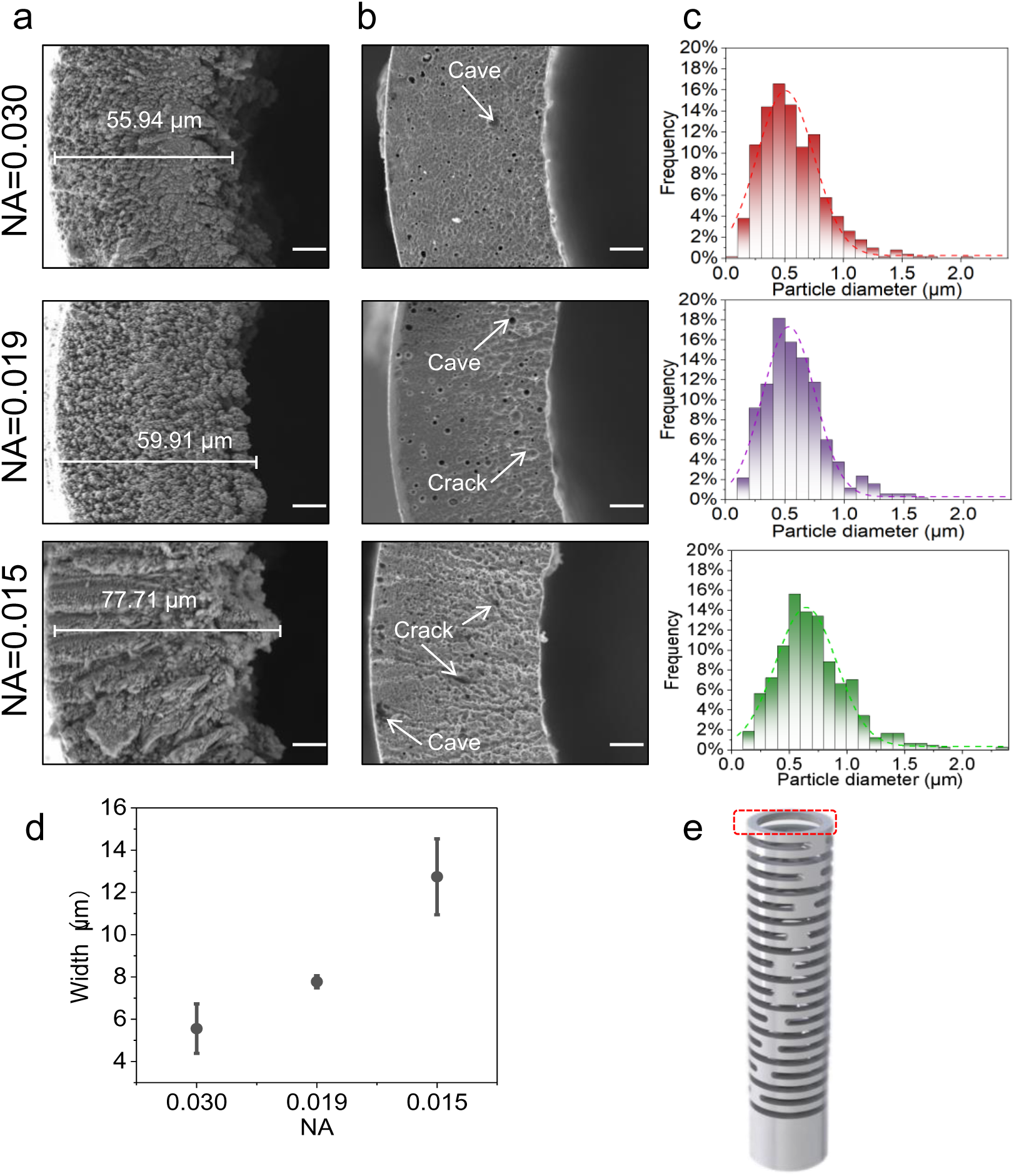
Aixal SEM images and quantitative analysis of the laser cut tube edges in Nitinol hypotubes fabricated with femtosecond laser. a, Axial SEM morphology of the fabrication hypotubes before acid dipping, NA=0.030 (top), NA=0.019 (middle), NA=0.015 (bottom). b, Axial SEM morphology of the fabrication hypotubes after acid dipping, NA=0.030 (top), NA=0.019 (middle), NA=0.015 (bottom). c, Frequency histograms illustrating the particle size distribution in axial SEM images of Nitinol hypotubes fabricated by femtosecond laser. d, Analysis of comparing debris width in axial SEM morphology at different NA, each group of NA is replicated three samples to minimize experimental variability. e, Schematic diagram of measuring the axial position. Scale bar, 10μm.

In Fig.2c, slight improvements in particle size distribution were observed as the NA was reduced from 0.030 to lower values. When transitioning from NA 0.030 to 0.019, the overall distribution of particles of different sizes remained relatively consistent, although there was a higher prevalence of particles smaller than 0.05 μm at NA 0.030. In contrast, upon changing the NA from 0.019 to 0.015, a noticeable increase in particles ranging from 1.5 to 2 μm was observed, suggesting that the use of a higher NA significantly contributed to the reduction in debris size.

Fig.2b illustrates the distinct surface topographies observed after the acid dipping process with varying NA values. With a reduction in NA from 0.030 to lower values, there is a noticeable trend where the axial view of the samples shows a greater amount of pits and grooves that are not only brittle but also prone to cracking and detachment. This indicates that the surface integrity is compromised at lower NA settings, leading to a less stable and more fragmented surface condition.

### Roughness analysis

Surface roughness was evaluated to scrutinize the impact of NA as an independent variable, and their associated effects on surface quality. Fig.3a and 3b displays the surface roughness data for various NA values. A comparatively minimal surface roughness of 8.07 μm was obtained at an elevated NA of 0.030. As the NA of the objective lens decreases, the accumulated debris height gradually increases, reaching 20.55 μm for the 0.019 NA lens and peaking at 31.63 μm for the 0.015 NA lens.

**Figure 3.**
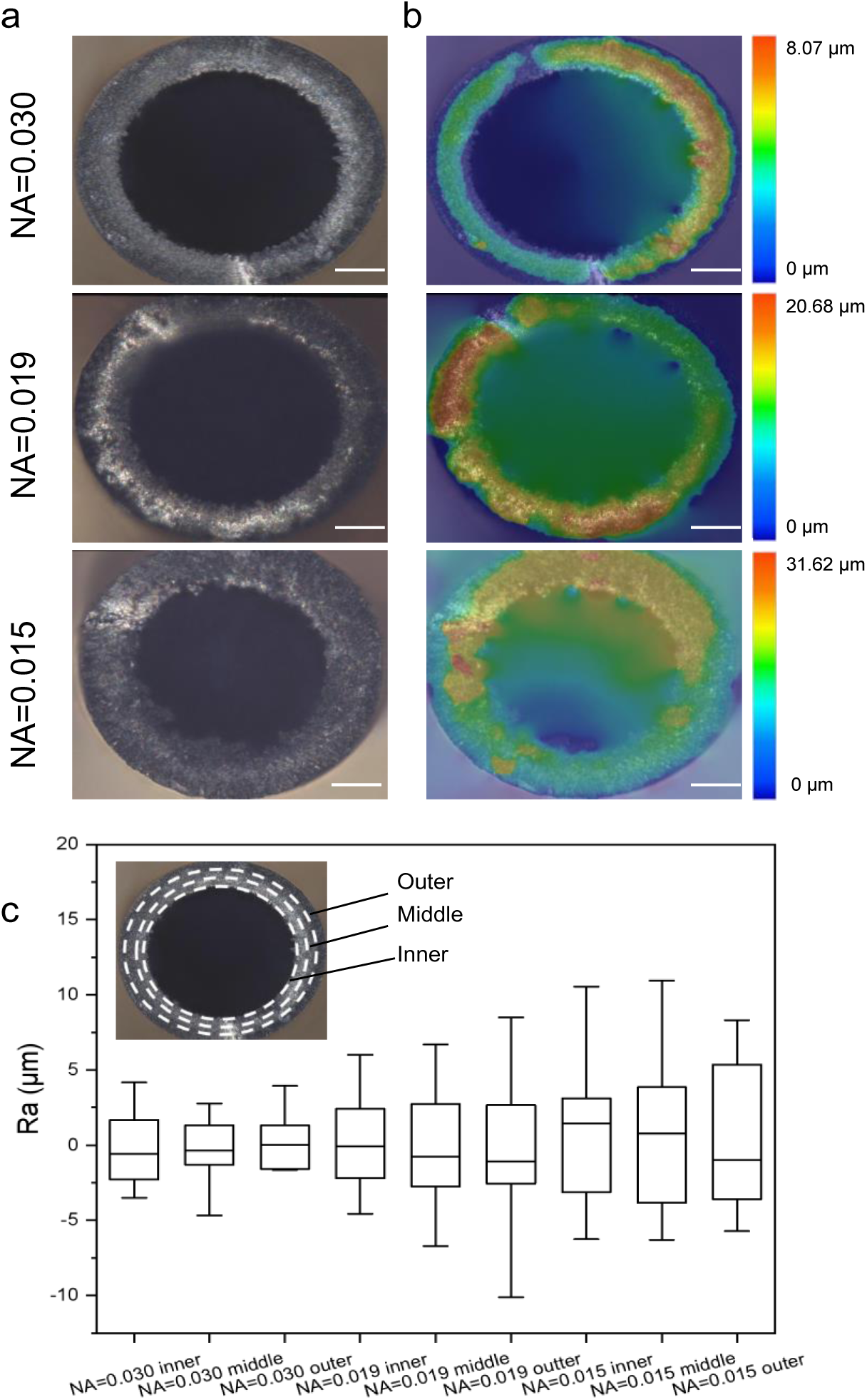
Nitinol hypotubes femtosecond laser fabricated surface roughness. a, Matrix of optical images, captured by a digital microscopy, shows the arithmetic mean height, and b surface roughness of the axial view fabricated by a femtosecond laser, NA=0.030 (top), NA=0.019 (middle), NA=0.015 (bottom). Scale bar, 50μm. c, A graph illustrating the line roughness comparison at varying NA, each group of NA is replicated three samples to minimize experimental variability. The top-left corner of the c, there are three white circles, which correspond to the line roughness on the inner, middle, and outer circles from the inside out.

Fig.3c presents the outcomes of the quantitative analysis conducted for the statistical evaluation of the independent variables and their influence on line roughness. An NA of 0.030 was demonstrated to have minimal variation in roughness from the inner to the middle and outer circles compared to all other NA-fabricated samples. When the NA was decreased from 0.030 to 0.019, there was little change in the inner roughness, but a noticeable increase in the middle and outer roughness was observed. As for the samples fabricated with an NA of 0.015, an increase in the surface roughness measured values from the inner circle to the outer circle was noted. The findings quantitatively revealed that the NA plays a higher contributing role in achieving optimal surface roughness.

### Dynamic mechanical analysis

Dynamic Mechanical Analysis (DMA) has been extensively employed to investigate the viscoelastic characteristics of composite materials in Fig.4a. The dynamic mechanical behavior of Nitinol hypotube is crucial because it indicates the mechanical properties of elastic materials across a broad temperature spectrum. Fig.4a illustrates the storage modulus. The storage modulus describes an approximation of the temperature-dependent load-bearing capacity, stiffness characteristics, and deformation resistance of the material. This study employs lenses of varying NA to produce Nitinol hypotubes with uniform mechanical structures. The Nitinol hypotubes created using high NA lenses with a value of 0.030 showed the highest storage modulus, suggesting that these high NA lenses enhance flexibility for clinical applications. Conversely, the hypotubes made with low NA lenses at 0.015 had the lowest storage modulus. Notably, the samples produced with medium NA lenses at 0.019 displayed a mid-range storage modulus. With increasing temperature, there was a consistent rise in the storage modulus values across all samples. Overall, it was found that higher NA were able to produce better elasticity.

**Figure 4.**
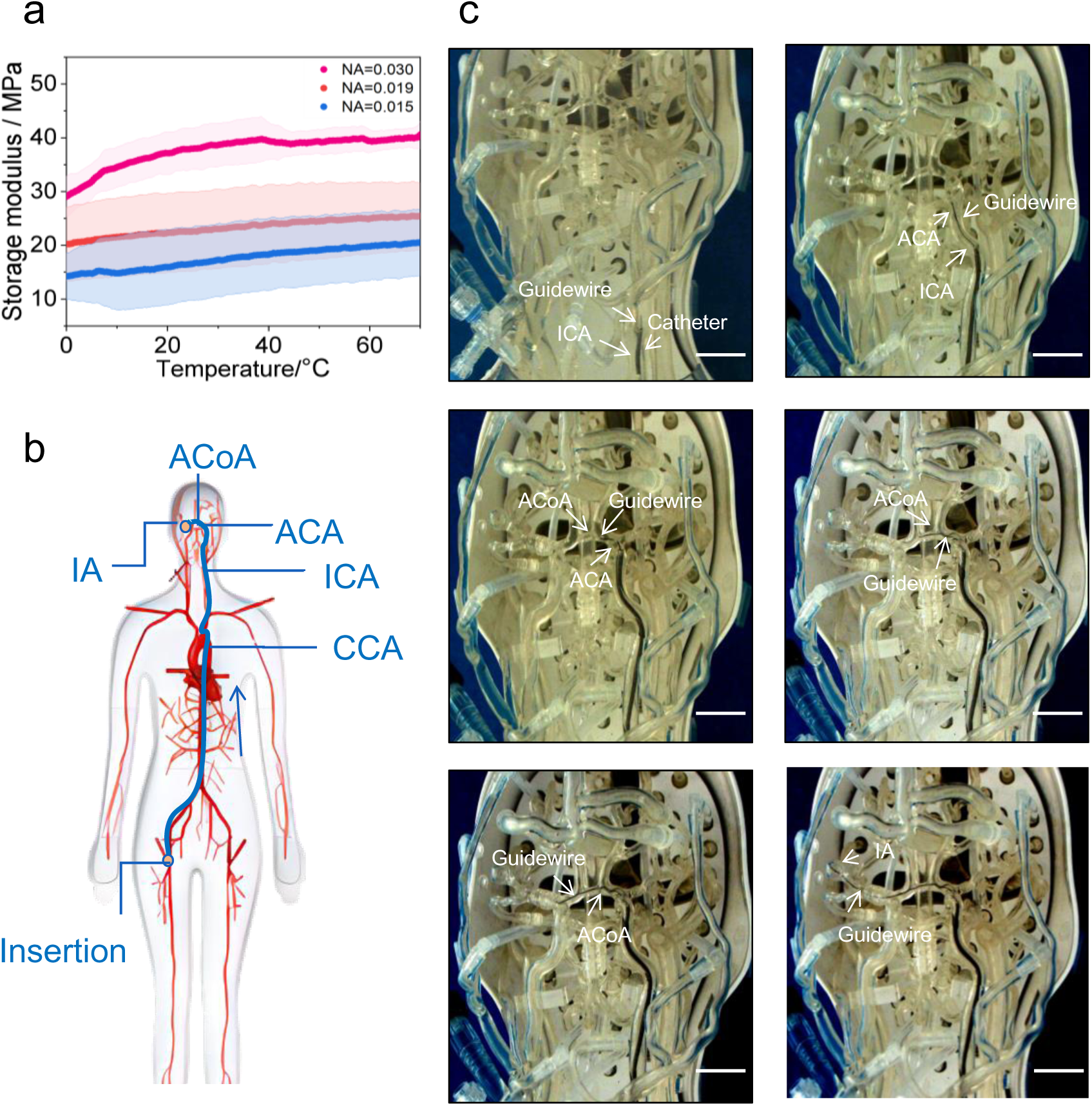
Mechanical capability examination of Nitinol hypotubes fabricated with femtosecond laser and in vitro experiment. a, Nitinol hypotubes DMA testing from 0 to 70 degrees Celsius, demonstrating elasticity for the specimen fabricated at varying NA, each group of NA is replicated three samples to minimize experimental variability. b, An overview of the human arteries vasculature with insertion point, travel paths, and target point. c, Nitinol guidewires made from the optimal storage modulus are used for in vitro experiments. The navigation along the arteries in a silicone model, starting from the femoral artery, into the ICA, across the ACA and ACoA to the IA. ICA, internal carotid artery;ACA,intracranial anterior cerebral artery; ICA, internal carotid artery; ACoA, anterior communicating artery; IA,intracranial aneurysm. Scale bar, 1 cm.

### Clinical performance analysis

The samples cut at an NA of 0.030 were subjected to mount, polishing, and ultimately fabricated into a neural guidewire to assess their potential clinical applications in surgical procedures for navigating complex anatomies. Figure 11a shows the path of a blood vessel navigated by a guidewire equipped with a Nitinol hypotube within a silicon model of the human vascular system. This experiment simulated a clinical trial where a guide catheter was first inserted into the brain, followed by navigation to the target site using a microcatheter. Figure 11b depicts a simulated patient with an intracranial aneurysm (IA).

Guidewires equipped with Nitinol hypotube fabricated were employed to navigate towards the anterior communicating artery (ACA) in cases of intracranial aneurysm (IA). The guidewire successfully passed through the sharp turn of the common carotid artery (CCA), then entered the internal carotid artery (ICA) at the neck, and finally navigated into the finer and more tortuous intracranial anterior cerebral artery (ACA) and anterior communicating artery (ACoA), reaching intracranial aneurysm (IA). Nitinol hypotube fabricated with a high NA (0.030) lens was fashioned into guidewires and subjected to in vitro testing. The results indicate that the Nitinol hypotube demonstrated excellent bifurcation and high steerability in the tortuous cerebral arteries. Overall, in vitro experiments demonstrate that guidewires fabricated using high NA lenses perform exceptionally well in practical applications, further supporting their feasibility for navigating complex in pathways.

### Finite Element Analysis

Finite element analysis was conducted to track the thermal distribution of the material expelled during the fabricating process. The thermal distribution imaging is depicted in Fig. 5a, featuring a concentric view of the tube as it is being cut. Data on thermal patterns and material removal were gathered to highlight variations associated with the independent variable of NA. The findings regarding the peak temperature experienced during fabricating across various NA settings are illustrated in Fig.a Employing an NA value of 0.030 resulted in a marginally elevated temperature in comparison to NA values of 0.019 and 0.015, with the temperature spike reaching 345K. Fig.5b indicates that at an NA of 0.030, the surface being machined exhibits reduced bulging and appears smoother. Additionally, a decrease in NA correlates with an increase in the number of bulges on the tube.

**Figure 5.**
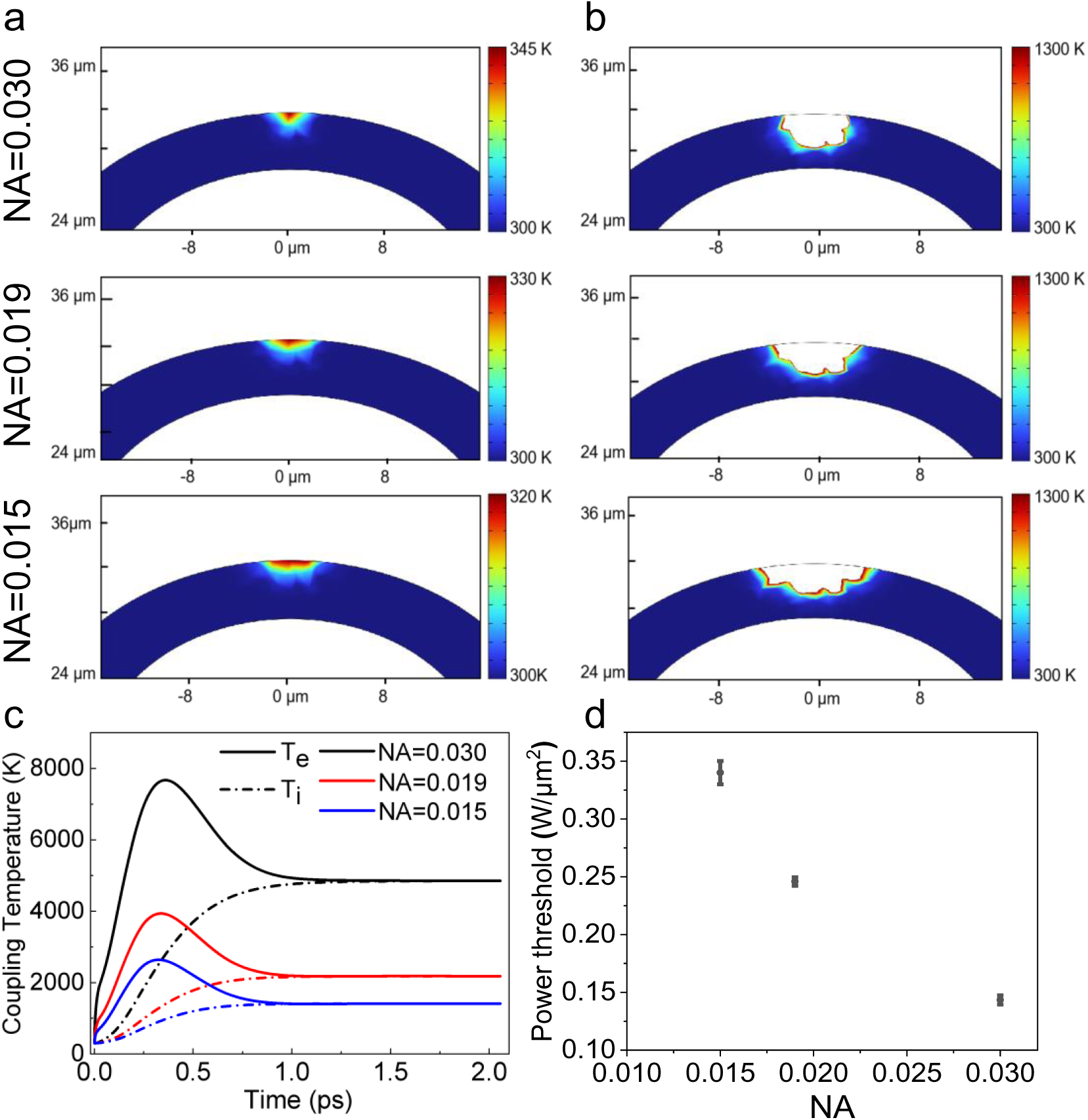
Two-temperature simulation on tube cross-section to demonstrate the difference in varying NA. **a**, The transient temperature distribution resulting from single-pulse energy, NA=0.030 (top), NA=0.019 (middle), NA=0.015 (bottom). **b**, The ablation status of the material with a NA of 0.030, 0.019 and 0.015. **c**, Electron and lattice temperature distribution in two-temperature simulation at varying NA, Te - electron temperature is solid line, Ti - lattice temperature is dashed line. **d**, Dependence of the threshold power for ablation kerf width on the NA of the objective lenses for femtosecond laser.

Fig.5c presents a comparison of the variations in electronic temperature and lattice temperature during fabrication with different NA values across a range of instances in the temporal domain. At an NA of 0.030, the instantaneous electronic temperature can approach 8000K, whereas, for NA values of 0.019 and 0.015, the temperatures are only 4000K and 2300K, respectively. Correspondingly, as illustrated in Fig.5d, an increase in NA results in a lower ablation threshold. The ablation thresholds for each NA were determined based on the Nitinol hypotube. For objective lenses with NA values of 0.030, 0.019, and 0.015, the respective ablation thresholds under these conditions were found to be 0.15, 0.26, and 0.34 W/μm². In conclusion, lenses with high NA generate a high energy density and localized temperature zones, highlighting their effectiveness in maintaining machining quality while managing thermal effects.

## Discussion

The initial qualitative analysis concerning the independent variable of NA provided an optimistic perspective on the potential benefits of femtosecond laser fabrication of Nitinol hypotubes with a higher NA. The qualitative observations indicated that the specimens fabricated with a higher NA laser demonstrated an enhanced integrity of the laser-fabricated surface compared to those manufactured with a lower NA. Fig.1 and 2 illustrates SEM imaging of the specimen fabricated with a high NA of 0.03, which showed minimal debris, contrasting with the specimens with NA values of 0.019 and 0.015, which exhibited a more substantial accumulation of debris. Additionally, the specimens fabricated with lower NA values presented a higher surface roughness due to the increased presence of resolidified material and pronounced striation patterns, in contrast to the specimen fabricated with NA=0.030. Subsequent sections will offer an in-depth quantitative discussion of the post-laser fabrication state of the Nitinol tubular test specimens, comparing the independent variables of NA.

Figures 1-6 and 9 illustrate that the kerf surfaces and axial surfaces machined with a high NA lens (0.030) exhibit superior surface quality. This is attributed to the ability of high NA lenses to focus the laser into a smaller area, generating higher energy density. This results in rapid material sublimation, reducing microcracks and surface defects^16^. In contrast, the lower NA lens (0.015), due to poorer focusing and lower energy density, results in higher surface roughness and more debris accumulation on the kerf surface^17^. The medium NA lens (0.019) yields intermediate energy density, displaying moderate surface quality and debris accumulation. Decreasing the NA of the objective lens results in a deterioration of the surface morphology^18^. With the increase in the NA, there is a concurrent decrease in the ablation threshold and fluence^19^. In the femtosecond laser fabricating of Nitinol materials under the influence of laser irradiation, Nitinol materials on the surface undergo direct vaporization. In contrast, samples fabricated by lenses with NA of 0.030 exhibits relatively higher optical fluence, facilitating the powerful ejection of debris. Hence, it was concluded that the High NA objective lens results in more localized material removal, with higher material ejection efficiency, which in turn improves the resultant surface integrity during Nitinol laser micromachining.

Additionally, the increased spot diameter caused by the lower NA results in a higher pulse overlap rate, which is another factor in the degradation of surface quality^20^. The increased pulse overlap rate leads to a higher overlap of thermal absorption, increasing kerf clogging. This explains the worsened surface morphology after acid dipping as seen in Figure 1c. The overlapping regions may not completely cool down or recover before subsequent pulses arrive, resulting in the accumulation of localized heating and ultimately leading to the formation of microcracks^21–23^. Therefore, the pulse overlap rate and the beam focusing effect due to the high NA are identified as critical factors that influence the final surface quality. A high NA leads to a reduced overlap rate, which can contribute to enhanced machining outcomes.

Figure 8 depicts the results of the storage modulus tests and temperature field distribution of Nitinol hypotube fabricated with different NA lenses. Samples fabricated with a high NA lens (0.030) exhibited the highest storage modulus, possibly attributed to the ability of high NA lenses to focus the laser beam into smaller areas, resulting in higher energy density^24^. This higher energy density reduces the temperature-affected zone of the material, leading to rapid sublimation and recrystallization on the material surface, in the case of higher NA, the laser beam achieves better focusing, resulting in enhanced energy concentration that improves fabricating precision and quality. This precise machining reduces material micro-cracks and defects, strengthening the spring’s storage modulus. Consequently, it mitigates internal stress accumulation within the material, thereby enhancing its mechanical properties^25^.

Furthermore, previous studies have revealed through metallographic observation that femtosecond laser machining does not alter the microstructure of nickel-titanium alloy materials^26^. This further confirms the high-quality machining capabilities of femtosecond lasers^13^. Interestingly, lenses with moderate NA exhibit variability in storage modulus and roughness. Specifically, a lens with NA=0.030 produces a smaller spot size and higher energy density, but lower overlap of laser spots. Conversely, NA=0.017 generates larger spots with lower energy density but greater overlap. For NA=0.019, the larger spot size and higher energy density result in an uneven energy distribution during machining. Specifically, at NA=0.019, although the spot size is larger^27^, the energy density is relatively higher, leading to concentrated energy in the overlapping regions between laser pulses. This uneven energy distribution causes irregular material removal, resulting in irregular surface and internal stress distribution, thereby affecting the stability of the storage modulus^28^. Compared to NA=0.030 and NA=0.017, the lens with NA=0.019 exhibits greater variability in surface roughness and storage modulus during sample machining.

In conclusion, the Nitinol hypotube fabricated with high NA lenses demonstrated excellent performance in vitro experiments, showcasing its superior mechanical properties and machining surface quality. The high energy density fabricating technology provided by high NA lenses reduces debris accumulation, maintaining the geometric precision and elasticity of the guidewire, allowing it to navigate through complex intracorporeal pathways smoothly. This result indicates that high NA lenses are an effective choice in femtosecond laser machining of Nitinol hypotube, capable of producing high-quality guidewires that meet the requirements of practical applications. This also provides important evidence for future practical applications and further research in Nitinol hypotube production.

## Method

### Materials

The experimental material consisted of a Nitinol tube characterized by an outer diameter of 330 micrometers and a wall thickness of 45 micrometers. The tube’s exterior surface featured a polished, non-oxidized finish, and the manufacturer (Peiertech) disclosed the inner surface condition of the tube. The compositional elements of the material were regulated in accordance with ASTM F-2063-18^29^, a standard that specifies the forged metallurgical criteria for nickel-titanium alloys employed in medical instrumentation. The elemental composition of the supplied Nitinol tubes reveals that the material is predominantly composed of nickel (Ni) at 54.49% by weight, followed by titanium (Ti) at 45.27%. Other elements present in smaller quantities include cobalt (Co), iron (Fe), and niobium (Nb) at 0.05%, 0.05%, and 0.025% respectively. Trace amounts of carbon (C), copper (Cu), chromium (Cr), hydrogen (H), nitrogen (N), and oxygen (O) are also detected at 0.04%, 0.01%, 0.01%, 0.005%, 0.005%, and 0.045% respectively.

### Characterization Procedures

Samples for testing were fabricated to a total length of 10 millimeters utilizing a femtosecond short-pulse laser. Two semi-circular removal fabrications were made around the tube using a laser to remove 80% of the material, thereby fabricating the specimen. The surface morphological analyses of the samples were carried out using scanning electron microscopy (SEM, ZEISS, Sigma 300) attached with Energy dispersive spectrograph (Oxford Instruments). All debris particle size and debris blockage severity were analyzed through Image J. After 10% nitric acid dipping through nitric acid, we employed SEM to observe the axial surface morphology and fabricate kerfs of Nitinol hypotube. The samples were positioned along their axis at the end for optical examination through a microscope (Z-100, Keyence) to analyze surface roughness. The presence of debris was assessed by quantifying the axial region of the tube fabricated by the laser; a larger area indicated a greater amount of debris on the sample. The surface roughness was determined by calculating the arithmetic mean height (Ra) over a circular area section of the Nitinol hypotube that was laser-fabricated. The microscope magnification factor of 700. The surface height profile was acquired using a Z-axis motor with a stepper resolution of 0.1μm. The dynamic mechanical properties of the samples were assessed using a DMA device (Mettler Toledo DMA1). These properties were evaluated over a 10 mm span, varying with temperature. The experimental temperature range was from 0 to 70 °C, with a heating rate of 2 °C/min. The dynamic strain was fixed at 1% with a frequency of 1 Hz. All quantitative analyses mentioned above were conducted with three replicates for each NA.

### Femtosecond laser micromachining setup

As shown in Supplementary Fig. 1, there is a depiction of the experimental setup employed to generate the femtosecond laser cut samples. The femtosecond laser micro-tube cutting setup depicted is a sophisticated system designed for high-precision cutting tasks. It includes a femtosecond laser as the primary light source, an optical shutter to control laser exposure, a half-wave plate for polarization adjustment, and a polarizing beam splitter to direct the beam. The function of a quarter-wave plate is to generate circularly polarized light, which is achieved by altering the polarization state of the incoming linearly polarized light. Meanwhile, a dichroic mirror is designed to reflect infrared femtosecond laser light, which is used for precise cutting, while allowing visible light to pass through for observation or imaging purposes. A CCD camera with illumination monitors the process, and an objective lens focuses the beam. The system features a micro-tube translation stage for the X-Axis, a rotary stage for Y-axis sample rotation, and a Z-Axis translation stage, all controlled by a dedicated software. An assisted gas nozzle is integrated for material processing assistance, and a motor control box drives the motion components, ensuring accurate and controlled cutting of micro-tubes. The specimens were created using Argon as the assist gas. A high-precision rotary motor, capable of a maximum rotational speed of 600 RPM, was affixed to a motion stage to facilitate the translation and rotation of the tube during the fabrication process. A femtosecond pulsed laser, with a maximum output power of 10 W,1030 nm wavelength, 240 fs pulse width, and a maximum repetition rate of 600 kHz, was employed for the fabrication of the specimens. The assist gas nozzle was aligned concentrically with the focusing optical assembly, allowing the laser beam to pass through the nozzle’s aperture. The material was secured in place beneath the nozzle with a hardened steel, high-precision bushing. The bushing was secured so that the center of the laser beam was positioned no more than 1 mm from the tip of the bushing. Once the setup was finished, the tube was cut along its central axis at the highest point using a single pass of the laser. The tube was rotated at different speeds while the laser power was applied, resulting in fabricating around the circumference at both ends of the tube.

### Design of experiments

The research sought to ascertain the distinctions among NA utilized in the femtosecond laser fabricating of Nitinol and to investigate the potential for achieving enhanced surface quality, mechanical properties, and clinical applicability. As shown in Supplementary Fig. 2, three air-spaced achromatic objective lenses with NA but similar structures have been designed. Three air-spaced achromatic objective lenses, each with a 3 mm pupil diameter but varying focal lengths—50 mm (NA=0.030), 77 mm (NA=0.019), and 100 mm (NA=0.015)—were employed to fabricate test specimens. The experimental setup involved three distinct NA, each processing three samples, totaling nine specimens. All nine specimens underwent analysis for debris, surface roughness, and DMA testing. Statistical analysis was conducted using Origin 2021 software.

The three guidewires with Nitinol hypotubes was maneuvered within a synthetic silicone model (Preclinc) containing the aorta, cerebral arteries, and major arteries of the lower and upper limbs. The filling liquid in the silicone vessel is a saline solution. This method is commonly utilized and is endorsed by the manufacturer of the vascular model for stimulating the clinical performance of guidewires^30^.

### Femtosecond laser ablation threshold

The Fraunhofer diffraction of a circular aperture can be employed to characterize a focused laser beam using a converging lens. When parallel light is vertically incident onto the aperture, which is positioned close to the front surface of the lens, Fraunhofer diffraction can be observed on the rear focal plane of the lens. Its diffraction aperture is *θ* (i.e. NA), definition of numerical aperture NA:

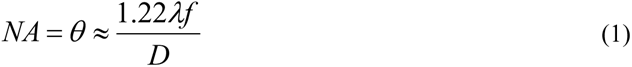

Calculating the material ablation threshold is essential for comprehending the process of femtosecond laser ablation of nitinol tubes. The ablation threshold signifies the minimum laser energy required to remove material from the working surface per unit area^31^. Here, *D* is the entrance pupil diameter, *f* is the objective focal length, and *W* represents the focused spot size, approximately equal to twice the Gaussian waist.

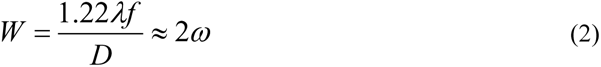

*D*_0_ represents the ablation kerf width. Its relationship with the power and ablation threshold of the laser can be expressed as^32,33^:

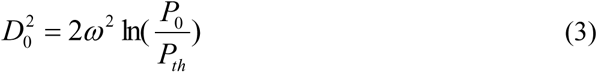

When the Power is constant, the relationship between the ablation threshold and the waist size of the Gaussian beam can be expressed as

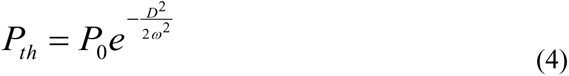

When the power of the femtosecond laser is constant, the effective pulse can be expressed as^34^

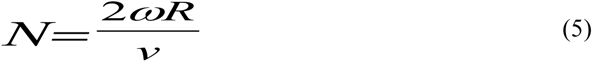

where *N* represents the effective number of pulses, while *v* is the rotating velocity. The spot size simulation was performed using Matlab 2020, and the modeling results are presented in the preceding sections.

### Finite element analysis for heat conduction

In this study, a three-dimensional cylindrical microtube model with a diameter of 330 μm, a thickness of 45 μm, and a length of 1 cm was constructed. The boundary conditions are established to determine the variation of parameters at the geometric boundaries with external variables within the computational domain. The initial temperature of the geometric model was set to 300 K, and the surface boundary conditions were zero flux (thermally insulated), maintaining a constant temperature of 300 K. Numerical solutions were obtained using an implicit solver based on the Backward Differentiation Formula (BDF) algorithm. At each time node, the temperature for all mesh nodes of the material was calculated. The time step was 1 fs, and the total solution time was 2 ps. The temperature-time and temperature-position variation curves for the electronic and lattice systems were deduced by fitting the numerical temperature values of computational steps and grid position nodes over the calculated time duration. We determined the values of *T_e_* and *T_l_* through the application of a pair of distinct equations.

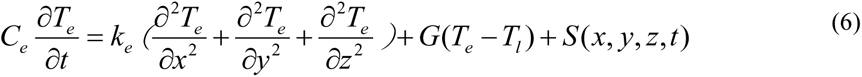

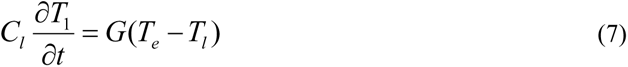

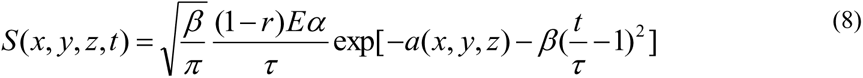

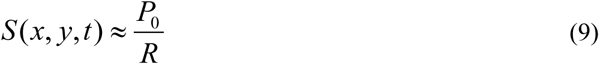

The electronic heat capacity is *C_e_*, the lattice heat capacity is *C_l_*, and the laser heat source term, *S* (*x*, *y*, *z*, *t*), are spatially and temporally distributed as Gaussian distributions. We use a single-pulse heat source. *G* represents the electron-lattice coupling coefficient. *β* = 4 ln 2 is constant. *τ* is the pulse width, *E* is the laser energy density, *α* is the laser absorption coefficient, and *r* is the laser reflectance. To model the independent temperatures of both electrons and lattice represented as *T_e_* and *T_l_* Here, NiTi electronic heat capacity is 67.5 J/m^3^/K^2^, electronic thermal conductivity is 28 W/m/K, and electron-lattice coupling coefficient is 46.44 × 10^16^ W/m^3^/K. Absorption and reflection coefficient is 4.22×10^7^/m and 0.65^35^.

## Acknowledgements

We would like to express our gratitude to YSL PHOTONICS and SUPER-WAVE for providing the femtosecond lasers.We are appreciative of YUNCO PRECISION for supplying the three-axis motors. We also thanks Z.Y. Li for the discussion about the genesis and progression of the project.

## Author Contributions

Y.F. Song and Z.W. Yang contributed equally to the manufacturing of the system. Y.F. Song was responsible for developing the image processing framework and analyzing all data, assisted by Z.W. Yang. G.X. and X.H. conducted the biological experiments, under the supervision of X.S. Yang. X.S. Yang drafted the manuscript, with editorial contributions from all authors. X.S. Yang oversaw all aspects of the project.

---

**Supplementary Note Figure 1.**
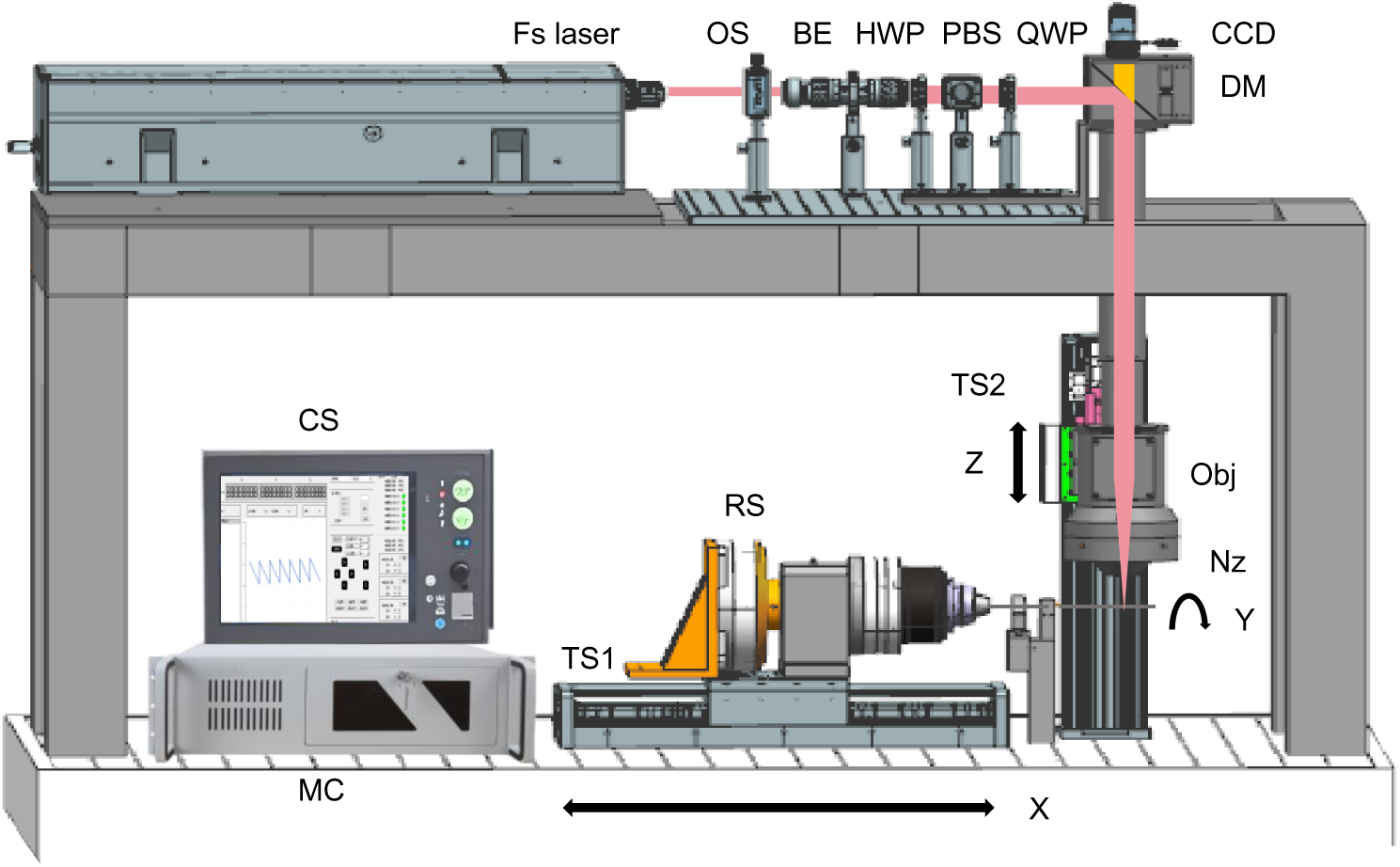
Femtosecond laser micro-tube cutting setup with three-axis motion components and control system. Fs laser,femtosecond laser. OS, optical shutter. HWP: half-wave plate.PBS: polarizing beam splitter. QWP, quarter-wave plate. DM, dichroic mirror. CCD, CCD camera with illumination.Obj,objective lens. TS1, micro-tube translation stage for X-Axis. RS, rotary stage for sample rotation on the Y-axis. TS2, micro-tube translation stage for Z-Axis. CS, control software. Nz, Assisted gas nozzle. MC, motor control box.

**Supplementary Note Figure 2.**
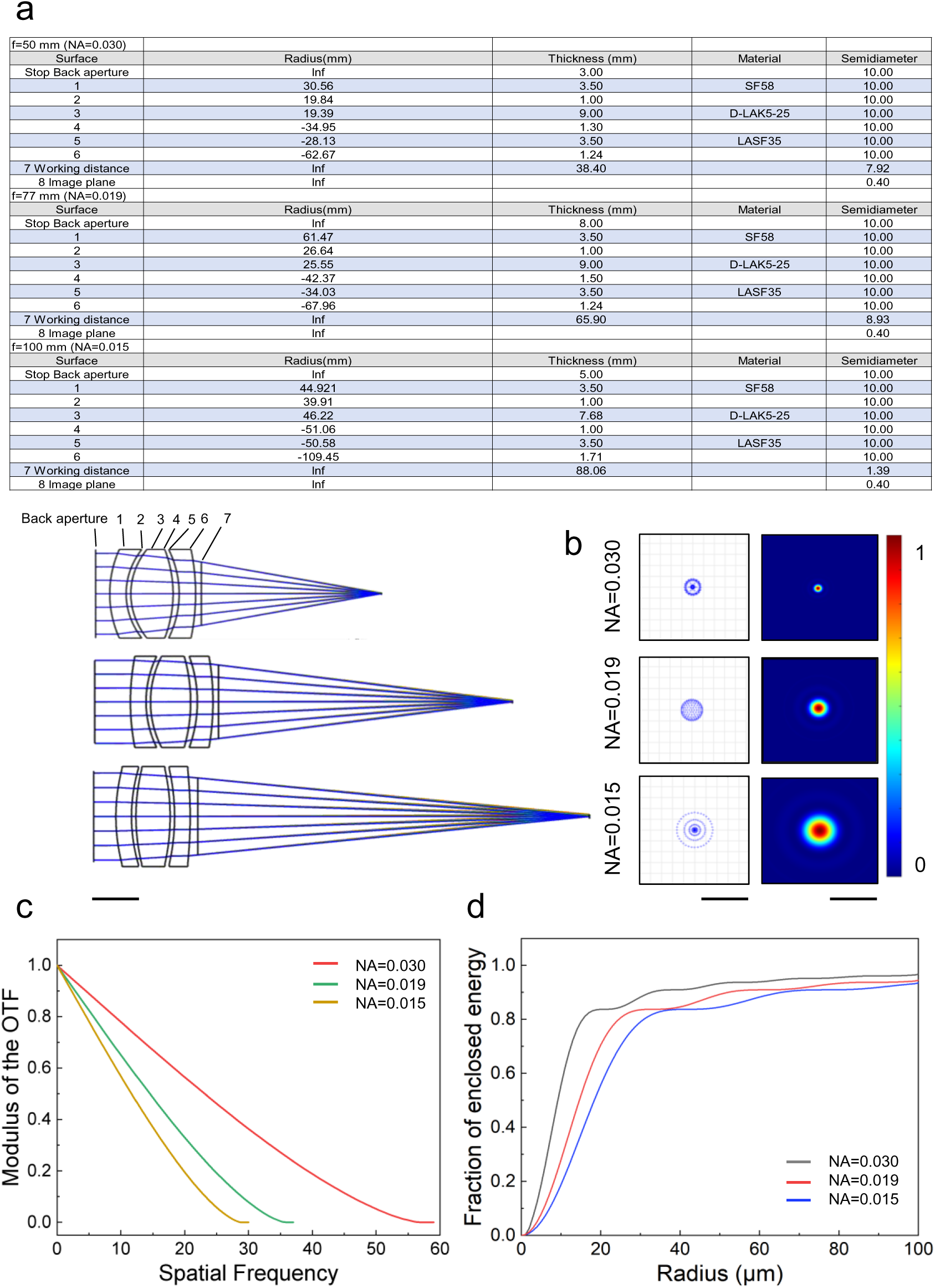
Optical design of femtosecond laser processing air-spaced objectives with different NA. a, The optical configuration that has been developed comprises six principal components to enhance its efficiency, the complete lens specification is furnished. The primary focus of the optimizations was centered on the 1030 nm wavelength for ultrafast laser processing.Scale bar, 10 mm. b, Left image is the spot diagram depicts the arrangement of light rays on the imaging plane that are directed towards the objective’s back aperture from a zero-degree angle, featuring a beam diameter of 3 mm and a wavelength of 1030 nm. Right image is the associated point spread functions emulated by Zemax. Scale bar, 100μm. c, The Modulation transfer function curves of different NA objectives featuring a beam diameter of 3 mm and a wavelength of 1030 nm. d, The Enclosed Energy curves of different NA objectives featuring a beam diameter of 3 mm and a wavelength of 1030 nm.

